# Improved peptide search for identification of SUMO and sequence-based modifications, in MaxSBM

**DOI:** 10.1101/2025.08.27.672604

**Authors:** Caroline Lennartsson, Pelagia Kyriakidou, Michael Lund Nielsen, Jesper Velgaard Olsen, Jürgen Cox, Ivo Alexander Hendriks

## Abstract

Post-translational modifications (PTMs), such as SUMOylation and ubiquitination, regulate key cellular processes by covalently attaching to lysine residues. While mass spectrometry allows site-specific identification of PTMs, most existing search engines are optimized for small, non-fragmenting modifications and struggle to detect large, fragmenting protein-based modifiers. We refer to these as Sequence-Based Modifiers (SBMs). To overcome this limitation, we developed an SBM-specific search strategy within MaxQuant that accounts for the fragmentation behavior of SBMs during peptide identification. Using publicly available datasets, we validated our approach for SUMO2/3. Our analysis identified distinct diagnostic features and characteristic mass shifts associated with SBM fragmentation, referred to in this study as d-ions (diagnostic ions) and p-ions. By leveraging these features, our method improved the identification of SUMOylated peptides from human cell lines by 13%, SUMOylation sites in mouse embryonic cells by 18%, and in mouse adipocytes by 25%. Our search method improved spectral annotation of SBMs by up to 9% increase in the median Andromeda score. Taken together, we highlight the potential of our SBM search to enhance the discovery of protein-based modifications.

**Highlights:**

- Development of a MaxQuant module tailored for identifying Sequence-Based Modifiers (SBMs), including SUMO2/3
- Incorporation of SBM-specific fragmentation patterns into search algorithms
- Enhanced biological discovery through improved PTM identification from mass spectrometry datasets

**In Brief:** Here, we introduce MaxSBM, an optimized framework for interpreting complex sequence-based modifiers (SBMs), particularly SUMO, within MaxQuant. Our approach incorporates SBM-specific d- and p-ion series into peptide scoring and annotation. By extending the theoretical spectral space to include fragments bearing partial SUMO (or other SBM) peptide remnants, MaxSBM provides a more comprehensive spectral annotation which enhances peptide scoring, resulting in increased identification rates at a higher confidence. Beyond methodological refinement, we validated MaxSBM via reanalysis of several physiological SUMO datasets, ultimately unlocking new insights via mapping of previously obscured modification sites.

## Introduction

One of the major strengths of mass spectrometry (MS)-based proteomics is the ability to identify post-translational modifications (PTMs). Identification of PTM sites is crucial for understanding biological mechanisms through direct identification of protein targets and regulatory dynamics. MS-based proteomics has become a powerful technique to study PTMs in a system-wide manner [1], [2]. While proteomics has advanced in identifying PTMs, detecting and pinpointing unstable (labile) modifications remains difficult, as they can degrade or be lost during analysis. Such modifications are, for example, SUMOylation and ubiquitination. In this paper, we present a new methodology for improved site-specific identification of these modification types.

Sequence-based modifiers (SBMs) are very important in biology. SBMs refer to protein-based modifiers that covalently attach to other proteins to regulate their function, stability, and various other molecular mechanisms. The earliest discovered SBM is ubiquitin, best known for its role in targeting proteins for degradation [3]. The family of ubiquitin-like modifiers has expanded over time, and includes SUMO1, SUMO2, SUMO3, NEDD8, ATG8, ATG12, URM1, UFM1, FAT10, and ISG15, each contributing to their unique functions [4]. Small Ubiquitin-like Modifiers (SUMOs) are a prominent subgroup of these modifiers that play critical roles in a wide range of cellular processes, and are particularly known for modifying the majority of nuclear proteins [5], [6]. SUMOs are covalently attached to lysine residues on target proteins via a conserved E1, E2, E3 enzymatic pathway [4], [7], [8]. Due to their high sequence similarity, SUMO2 and SUMO3 are considered biologically dispensable [9], while SUMO1 has different roles [10]. Conjugation of SUMO2/3 contributes to many essential cellular mechanisms; nucleocytoplasmic exchange [11], DNA damage response [12]–[14], mRNA processing and metabolism [15], chromosome biology and chromatin organization [16]–[18], cell differentiation [19], gene transcription regulation [16], [17], [20]–[22], and far more. Consequently, the modification is also observed in many pathophysiological processes such as tumorigenesis [23], diabetes [24], heart failure [25], [26], neurological disorders [27], and liver diseases [28]. Understanding SUMOylation by mapping of the subset of proteins that are SUMOylated (i.e., the SUMOylome) is therefore crucial to elucidating key parts of cellular physiology.

The main characterization of SUMOylation sites in a systems-wide manner is done via mass spectrometry-based proteomics [5], [6]. Endogenous SUMOylation identification is done with an extended MS workflow, involving antibody-based enrichment and enzyme serial digestion. This workflow generates a sequence remnant on the target peptides, which is difficult to identify due to its complex fragmentation pattern [5]. MS computational workflows begin with centroiding and deisotoping of the raw spectral data. The cleaned spectra are searched against theoretical peptide spectra derived from in silico digestion of a protein database. Peptide-spectrum matches (PSMs) are made by comparing experimental spectra to the theoretical ones using search engines, such as Andromeda [29] (used in MaxQuant [30]) or MSFragger [31]. The Andromeda search engine employs a probabilistic scoring algorithm to identify peptide-spectrum matches (PSMs). PTM-containing peptides are identified, and the PTM itself is localized within the peptide by finding fragment ions bearing mass deltas that correspond to the mass of the modification. Andromeda evaluates a potential modification neutral loss from a modification by calculating the scores, both with and without the loss. The higher of the two scores is retained as the final score [29]. To ensure statistical robustness, all scoring steps are mirrored identically against a reversed decoy database for false discovery rate (FDR) estimation [29].

There are many challenges related to the identification of SBMs, such as SUMOylation. They are difficult to study due to being biologically transient or low-abundant, and notably generate convoluted fragmentation patterns during MS analysis. Some progress has been made to improve their detection; for instance, MSFragger [31] can identify complex PTMs in two modes. Firstly, the tool PTM-Shepherd can search for diagnostic features of fragmenting PTMs [32]. Secondly, MSFragger-Labile can increase labile PTM identification through consideration of PTM remainder masses on the peptide ions [33]. These advancements have been shown to enhance identification of glycosylation and ADP-ribosylation [32]–[34]. The software pLink [35] and OpenUaa [36] were developed for analysis of crosslinked peptides, and have previously been used for identification of SUMOylation because SUMO-modified peptides consist of two covalently linked sequences, a scenario similar to crosslinked peptides in MS/MS analysis. However, the currently available versions of this software no longer support SUMO-specific searches. Finally, the SUMmOn software was developed to identify SUMOylation [37]. SUMmOn utilizes an initial database search with SEQUEST[38] and then looks for SUMO peak patterns. However, the software has only seen limited use in practice [39] and depends on software compilers that are not compatible with contemporary computer hardware. Taken together, the efforts made towards identification of SBMs and labile modifications highlight the overall interest and necessity for such approaches, and a need for optimized and up-to-date software still exists.

The major challenge to overcome is the analysis of convoluted MS/MS spectra resulting from the fragmentation of peptides bearing SBMs. Because the SBMs themselves fragment, the result is a much more complex fragmentation pattern that involves three peptide termini, rather than two. The SBM ion series integrates with the y and b ion series of the target peptide, hindering identification of the target peptide sequence. Here, we anticipate, evaluate, and incorporate series of fragment ions and remainder masses into the theoretical search space for peptide-spectrum matching (PSM) into a module in the MaxQuant search engine [30], [40]. Since the strategy is to expand the theoretical search space, we must account for an increased probability of getting decoy matches. This problem is widely described in the database search field [41]–[43], especially in proteogenomics [44].

To address these challenges, we designed a dedicated MaxQuant module specifically for SBM identification. This module annotated and enhanced the identification of SBMs during a dedicated data-dependent acquisition (DDA) database search. To optimize and evaluate our SBM module, we utilized several representative published data sets based on endogenous SUMOylation: HEK cell lines [5], [19], [45]. With the MaxSBM module, we improved the identification of SUMOylated peptides in HEK cell lines by 13%. Additionally, we observed an increase in detected SUMOylation sites in mouse samples: 18% in embryonic cells and 25% in adipocytes. The method improved the spectra annotation of SBMs by higher PSM scoring and better overall intensity and peak coverage. We also compared our software with the existing tool for labile PTMs, MSFragger-Labile, and with a comparable pipeline, using Percolator for significance estimation, we showed that MaxSBM yields a higher identification rate. Collectively, these findings highlight the utility of our approach for the identification SBMs in proteomics, thereby enabling deeper biological insights and advancing our understanding of the functional roles of these PTMs in health and disease.

## Experimental Procedure

### Experimental Design

#### Datasets

We used four published proteomics datasets to develop and evaluate the SBM module. For clarity, each dataset is assigned a shorthand name that will be used consistently throughout the manuscript. We analysed three SUMOylation and one ubiquitination datasets: SUMO-HEK, SUMO-Adip, SUMO-MEC, and Ub-LysC.

The SUMO-HEK dataset (PXD008003) published by Hendriks et al. [5] was derived from HEK cells exposed to proteotoxic stress (heat shock). It was generated using a site-specific proteomics strategy that combines a dual-protease digestion method with peptide-level immunoprecipitation to map endogenous SUMO2/3 modification sites. Specifically, denatured lysates are first digested with Lys-C, peptides enriched via the SUMO2/3 targeting antibody, and then further digested with Asp-N before LC–MS/MS analysis (Fig. 1A).

**Figure 1.**
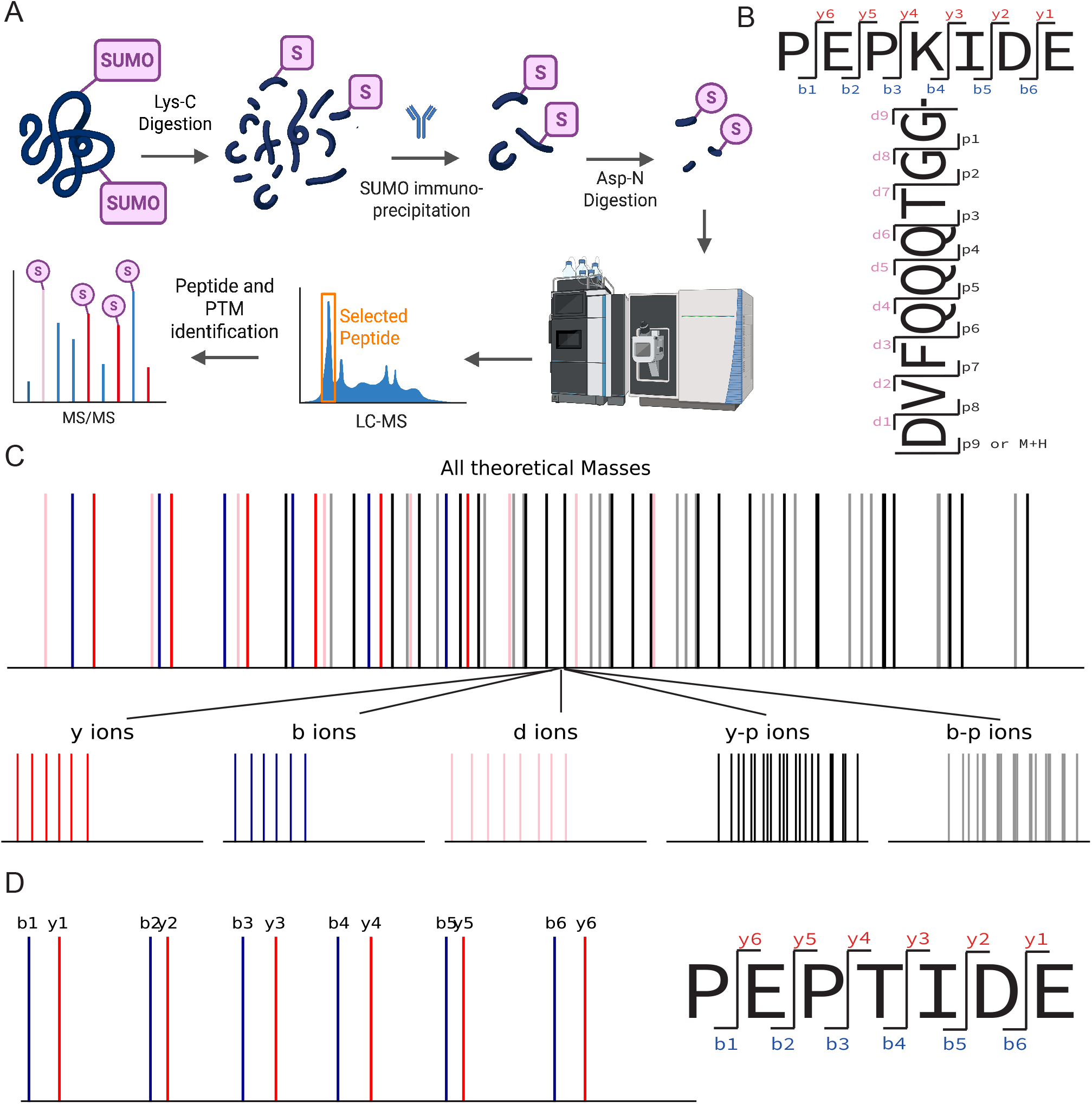
A. General method workflow for MS-based identification of endogenously SUMOylated peptides [5]. The samples are digested with AspN and LysC enzymes. The SUMOylated peptides are enriched with antibody-based enrichment. B. SUMOylated target peptide displaying the y-, b-, d-, and p-ion series. The y- and bions can potentially bear a sequence-based modifier (SBM) fragment (p-ion) and are annotated with both their corresponding y-/b- and p-ion labels. The diagnostic (d-ion) series, representing SBM fragments that are no longer attached to the target peptide, is shown in pink. C. Theoretical spectrum showing all possible fragments for the SUMOylated peptide depicted in panel B. The theoretical spectra consist of y-, b-, d-, y-p, and b-p -ion series. D. Theoretical spectrum of a standard (unmodified) peptide, shown for comparison to highlight the increased fragmentation complexity introduced by SUMOylation.

The SUMO-Adip dataset (PXD024144), described by Zhao et al. [45], was generated to profile SUMOylation dynamics during mouse adipocyte differentiation. SUMO-modified peptides were enriched with a SUMO2/3-specific antibody and analysed by LC–MS/MS to map stage-specific SUMO targets across differentiation.

The SUMO-MEC dataset, described by Theurillat et al. [19] (PXD017697), compares SUMOylomes between mouse embryonic stem cells and fibroblasts, highlighting developmental specificity.

The Ub-LysC dataset, reported by Akimov et al. [46] (PXD006201), was generated using a unique ubiquitination detection method based on Lys-C digestion, which leaves a longer covalently attached remnant with the sequence ESTLHLVLRLRGG (1431.83 Da).

#### Search Settings

##### MaxQuant

##### SUMO datasets (HEK, Adip, MEC)

All SUMO datasets were searched in MaxQuant (version 2.6.8.0) with default parameters unless specified. To accommodate SUMO remnants, the maximum peptide mass was set to 6000 Da, the second-peptide search was disabled, the maximum number of variable modifications per peptide was set to three, sequence-based modifier was enabled, and up to eight missed cleavages were allowed. Enzyme specificity used AspN, LysC/P, and GluN. The fixed modification was set to Carbamidomethyl (C) and the variable modifications to Oxidation (M) and Acetyl (Protein N-term). SUMO was included as an additional variable modification using our SBM variants (Standard SUMO, SBM 0, SBM 3, and SBM 8), depending on each run, with d and p-ions selected from Table 1.

**Table 1.**
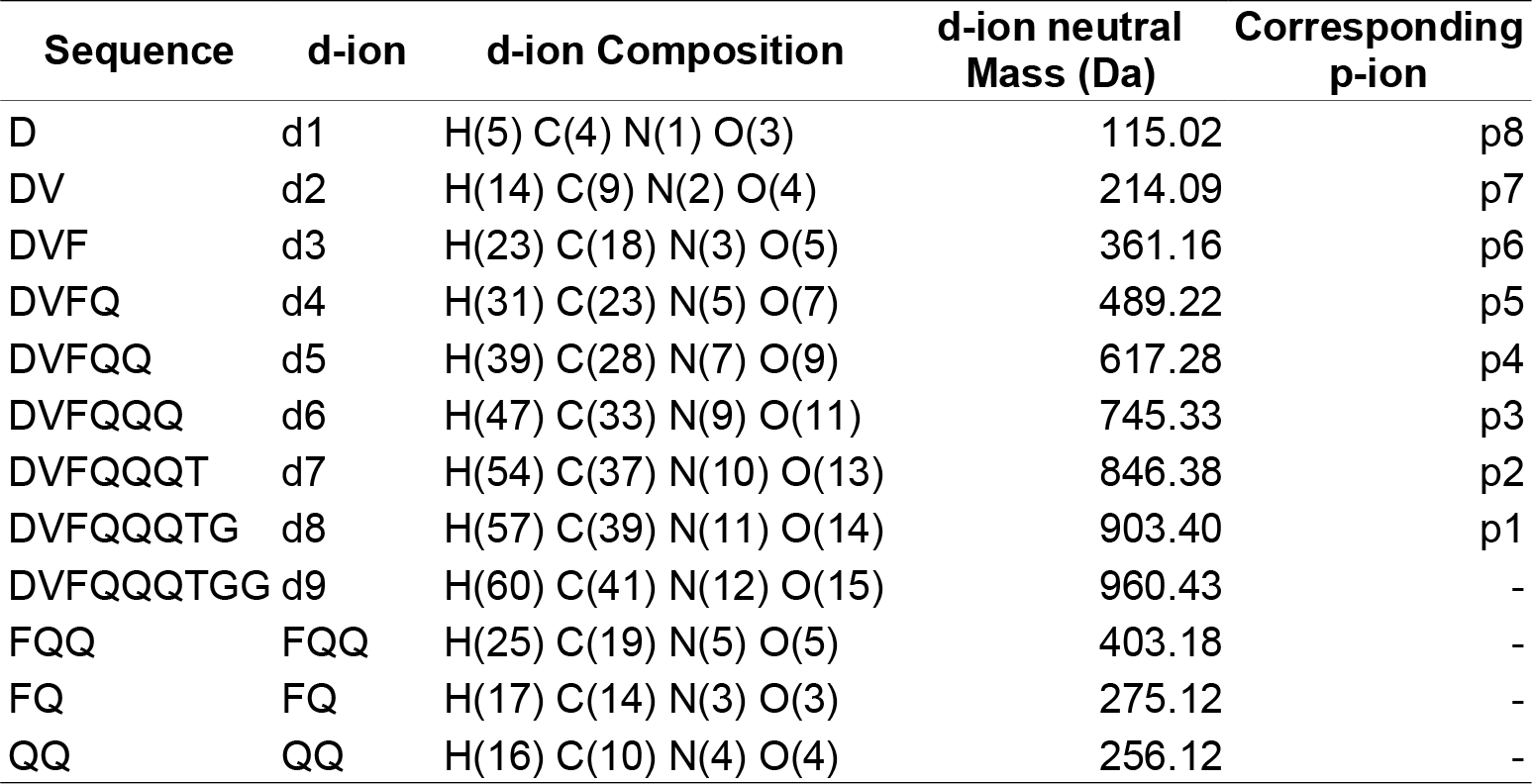
Chemical compositions and monoisotopic neutral masses of SUMO d-ions and corresponding p-ions, including doubly fragmented sequences FQ, FQQ, and QQ.

##### Optimization runs (SUMO)

Before the main analyses, we ran small optimization searches on a subset of SUMO-HEK raw files to tune the SBM search space efficiently. We tested different counts and selections of p-ions and evaluated alternative sets of diagnostic d-ions. These runs were used only to choose parameters for the main analyses. The optimization runs and the subsequent main HEK searches used the same UniProt [47] human proteome FASTA (downloaded December 2020, 97,795 entries including isoforms).

##### SUMO-HEK

HEK searches used the UniProt [47] human proteome (downloaded December 2020, 97,795 entries including isoforms). We performed four searches: Standard SUMO, SBM 0, SBM 3, and SBM 8. We also performed a dedicated MaxQuant run to get the input for Percolator that was identical to SBM 3 except for the internal filtering parameters, which were set as peptideFdr, proteinFdr, and siteFdr set to 1 (100%), minScoreModifiedPeptides, and minDeltaScoreModifiedPeptides set to 0 to produce an unfiltered msms.txt for downstream rescoring in Percolator.

##### SUMO-Adip and SUMO-MEC

These searches used the UniProt [47] mouse proteome (downloaded March 2022, 63,654 entries including isoforms). For each of these datasets, we performed two searches: a Standard SUMO search and an SBM3 search.

##### Ubiquitin group (Ub-LysC)

Ubiquitin searches were performed in MaxQuant version 2.6.2.0. Settings matched the SUMO runs except for the following differences. The maximum peptide mass was set to 6500 Da, and enzyme specificity used LysC/P rather than the combined AspN, LysC/P, and GluN rule. Ubiquitination was chosen as an additional variable modification using the remnant ESTLHLVLRLRGG (1431.83 Da), with d and p-ions selected from Table S1. Ubiquitin searches used the UniProt [47] human proteome (downloaded September 2021, 100129 entries including isoforms).

##### Percolator

To compare MaxQuant and FragPipe/MSFragger fairly, we rescored the MaxQuant PSMs with Percolator (version 3.06) using the Crux toolkit (version 4.2) [48], [49]. MaxQuant assigns each PSM a posterior error probability-based score (PEP score) to estimate the chance that a given PSM is incorrect [40]. FragPipe uses Percolator, which trains a support vector machine (SVM) on features derived from searches against target and decoy sequences [50]. By running MaxQuant results through Percolator as well, both pipelines use the same PSM-level rescoring step for false discovery rate estimation. This makes the comparison more fair because differences mainly reflect the search settings and engines, not different PSM level rescoring methods. We note that the feature sets supplied to Percolator differ across setups. The Percolator standalone run based on the MaxQuant msms.txt uses a feature set that is not identical to the one passed to Percolator inside FragPipe.

As described in the MaxQuant SUMO-HEK section, we generated an SBM 3 export run to produce an unfiltered msms.txt for Percolator. In that export run, the MaxQuant FDR and some score thresholds were switched off to remove any filtering before rescoring.

Percolator is typically used with inputs that provide separate evidence for the target and the decoy for each spectrum, a best target score, and a best decoy score. MaxQuant, in contrast, reports only the highest score overall after searching against both target and decoy sequences, so msms.txt contains one score per spectrum that can correspond to either a target or a decoy. Following community practice reported by others [51], we used msms.txt as Percolator input despite this difference. To do so, we reformatted the table to Percolator’s expected column layout and names and assigned the appropriate target or decoy label to each PSM without constructing explicit target–decoy score pairs. The feature mapping used in this conversion is provided in Table S2, and the conversion script is available at https://github.com/carolinelennartsson/SUMOylation. Percolator was executed with default settings except that the --only-psms flag was set to True to report on PSM level. Percolator uses three-fold cross-validation for SVM training [52].

##### MSFragger

We also processed the data with the MSFragger-Labile search (MSFragger version 4.1) [33] in the FragPipe MS analysis pipeline (version 22.0) to provide a comparison with MaxQuant. Statistical estimation of the MSFragger-Labile search output was calculated with Percolator (version 3.06.5) [50]. The decoys and contaminants were added to the FASTA sequence database with Philosopher [53]. Enzymes were set to LysC/P and a custom rule that cleaves N-terminal to D and E (to mirror the AspN/GluN behavior used in MaxQuant). SUMO was set a variable modification with mass 960.43 Da, on lysine residues (K). The diagnostic ions that were included in the search were d2 to d8 (Table 1). The remainder fragment ions included were selected from p1 to p8 (Table 1), with either 0, 3, or 8 number of p-ions included. When three remainder ions were included in the search, p2, p3, and p7 were selected. PSMs were filtered at 1% FDR for all searches, as per default.

#### Post identification processing and data analysis

All post identification processing was done in Python using pandas [54] and numpy [55]. The results were visualized with matplotlib [56] and seaborn [57]. The PSM data was extracted from the *‘evidence.txt’, ‘msms.txt’*, and *‘sites.txt’* MaxQuant output files. For analysis of the MaxSBM output, the modification table *‘sites.txt’* is recommended. Reverse and contaminant hits were removed from the result when they were not explicitly visualized. Additionally, PSMs containing two or more SBMs were filtered out. To benchmark the number of detected gene names, the peptide to protein inference was done by finding all matching proteins for each peptide sequence from the protein database. A representative protein was chosen for each PSM through canonization. This was done by selecting the protein with the best annotation from the UniProt reference proteome [47]. The supporting data for the FASTA files were downloaded from the UniProt web server. A protein was selected from the protein group by its annotation in the UniProt database in the order: review status (reviewed or unreviewed), annotation (0-5), number of Gene Ontology (GO) terms, alphabetical order of entry name, and finally entry name length (older entries have shorter names). During protein inference, the SUMO modification was only permitted at the C-terminus of a peptide if it was followed by an E or D in the protein sequence. The Python package Biotite was used to read and parse the FASTA files [58]. Structural analysis of the SUMO sites was done with the AlphaFold predicted database of protein structures [59], [60]. The AlphaFold database did not contain structures of the isoforms, which were therefore omitted in the analysis. Pairwise comparisons between the variable modification search (VM) and the other searches were performed using unpaired two-sample t-tests with unequal variance assumptions, implemented via the ttest_ind function from the scipy.stats Python library [61]. This analysis was applied independently to intensity coverage, peak coverage, and the number of matched peaks. P-values from these tests were annotated on the corresponding violin plots in scientific notation alongside their significance levels. Significance thresholds were defined as follows: p□<□0.05 (*), p□<□0.01 (**), p□<□0.001 (***), and p□<□0.0001 (****); values equal to or above 0.05 were considered not significant (ns). All scripts for visualization are available in the GitHub repository: https://github.com/carolinelennartsson/SUMOylation.

### Statistical rationale

No new experiments were conducted in this study. All analyses were based on previously published datasets and leveraged established statistical frameworks as described in earlier studies.

## Result

### Implementation of MaxSBM

We developed MaxSBM, a module integrated into MaxQuant, with the aim of enhancing the identification, annotation, and validation of SUMOylation sites. To this end, we leveraged previously published endogenous SUMOylation MS datasets from human HEK cells (SUMO-HEK) [5], mouse embryonic cells (SUMO-MEC) [19], and mouse adipocytes (SUMO-Adip) [45]. The protocol for identification of endogenous SUMOylations sites includes; serial enzymatic digestion, antibody-based enrichment, and MS-based peptide identification, The method is described in detail by Hendriks et al. (2018) [5], summarized is shown in Fig. 1A. Digestion using both Lys-C and Asp-N of target proteins and the covalently attached SUMO2/3 protein, leaves a subset of peptides bearing the SUMO2/3-derived sequence remnant (DVFQQQTGG) on lysine residues (Fig. 1B). The theoretical combinations of single and double fragmentations of SUMOylated peptides are highly convoluted (Fig. 1C), and include the standard b- and y-ion series, as well as two additional ion series specific to SBM modification: the diagnostic fragment ion series (d-ions) and the modification loss series (referred to here as p-ions) (Table 1 and Fig 1B). Compared to the theoretical spectra of an unmodified peptide (Fig. 1D), the y-p and b-p -ion peaks substantially increase the number of theoretical peaks (Fig. 1C). To consider this, we developed a new module MaxSBM that builds on top of Andromeda by incorporating the SBM ion series into the search space (Fig. 2A).

**Figure 2.**
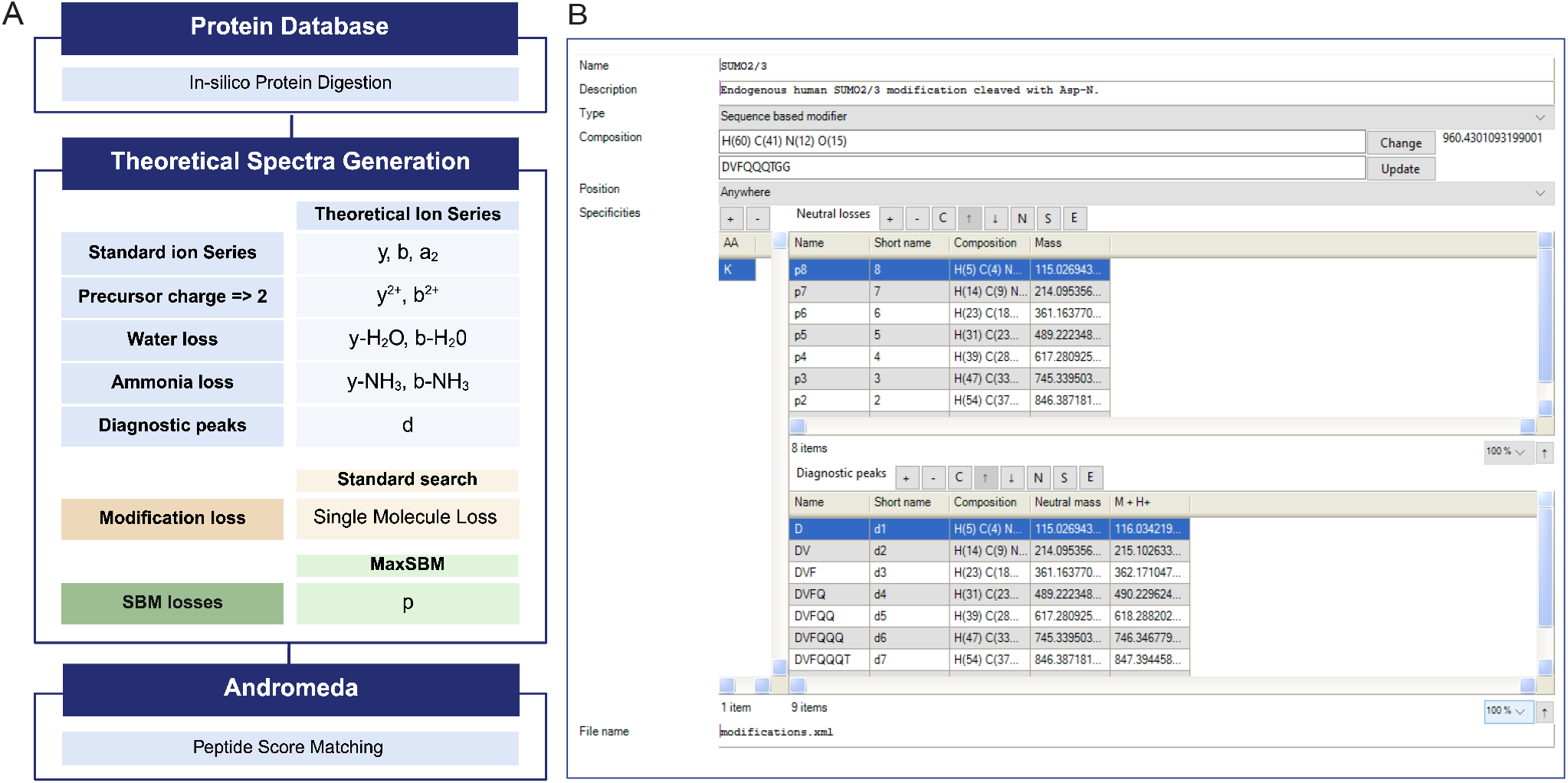
A. Theoretical Peak List Generation for Peptide Score Matching in Andromeda [29]. During peptide score matching, a protein database is in-silico digested into peptides. A theoretical peak list is generated for each peptide for the possible ion series. In the standard module, only a single modification loss is considered, whereas in the SBM module, all specified losses are utilized for identification. B. MaxQuant graphical user interface for defining SBM modifications, which allows users to define SBM modifications by inputting an amino acid sequence. The program then automates the process by calculating theoretical losses for SBM, where p-ions are displayed under “Neutral Losses” and the d-ions under “Diagnostic Peaks.”

### MaxQuant GUI configuration

To enable the detection of SUMOylated peptides using MaxQuant, we configured the software to support sequence-based modifications (SBMs) via the “MaxSBM” module. These settings were implemented through the graphical user interface (GUI) in the modification list, under the ‘*configuration’* tab (Fig. 2B). The MaxSBM is activated by incorporating a variable modification in a DDA search with the modification type *‘sequence based modifier’* (Fig. 2B). For SBMs, masses are calculated automatically when prompted a sequence input in the ‘*sequence’* box. After the search is finished, the identified p-ions are shown in the MaxQuant spectral viewer for the identified SUMOylated spectra (Fig. S1A and B). The identified p-ions are reported with annotation and mass in the *‘msms.txt’* search output table. The MaxSBM output with the complete set of modification identifications are found in the modification output table *‘sites.txt’*. We recommend analysis with the downstream proteomics analysis platform Perseus [62].

### Optimization of the search space for efficient identification of SUMOylation

To ensure optimal performance of the new SBM module, we systematically optimized the theoretical search space by evaluating the impact of p-ion inclusions. All optimizations were done on the endogenous SUMOylation dataset (SUMO-HEK) [5]. Although adding p-ions to the theoretical search space increases the possibility of identifying target peptides, the increased number of theoretical peaks also raises the chance of a decoy match [29]. Therefore, careful tuning was necessary to strike a balance between sensitivity and specificity. We optimized on the HEK cell line dataset, as it’s the most extensive identification of human SUMO sites [5]. To fine-tune, we focused on two key aspects: identifying which p-ions yield the highest number of SBM identifications, and determining the optimal number of p-ions to include. Firstly, we found that while there was only a slight preference in the benefit of each, p2, p3, and p7 were optimal to include for getting the highest identification rates (Fig. S2A). Secondly, we performed a series of searches wherein we searched a combination of 2, 3, 4, 5, 6, and 8 p-ions, preferentially selecting those that individually gave the highest identification rates. We found that with the HEK dataset, the optimum number of SUMO p-ions to include was three (Fig. S2B). In addition to optimizing the p-ions, we also tested the effect of the diagnostic (d-ion) series on the identification rate. To show the prevalence of the d-ions across the dataset, we summed the d-ion peaks from the SBM 3 search. We observed, as expected, that the d-ion series was frequently observed across the measure peaks (Fig. S3A). For analysis of the d-ions influence on the identification rate, we compared searches including three d-ion settings: none of the d-ions, d2 to d9 and an optimized set, reported by Hendriks et al. [5]. The optimized set included d2-d9, along with masses corresponding to double-fragmented d-ions with the amino acid sequences ‘FQQ’, ‘QQ’, and ‘FQ’ (Table 1.). Regardless of their prevalence, we showed that the number of d-ions did not influence the identification rate (Fig. S3B). Nonetheless, including more d-ions increased spectral peak coverage (Fig. S3C), the number of matched peaks (Fig. S3D), as well as the spectral intensity coverage (Fig. S3E). Thus, while d-ions can increased the quality of spectral annotation and served as an internal validation for the presence of SUMOylation, they otherwise do not positively or negatively affect identification rates.

### Enhanced identification rates using MaxSBM

We leveraged a previously published endogenous SUMOylation MS dataset from human HEK cells (SUMO-HEK) [5] to enhance identification rates using MaxSBM. With the SUMO-HEK datasets, we showed that including the SBM peaks in the search improves the annotation of SUMOylated peptides (Fig. 3). We evaluated the performance of the SBM module under different configurations of p-ion inclusion. We tested different p-ion configurations, comparing a variable modification (VM) search without p-ions, a standard MaxQuant search with one neutral loss, the optimal three-p-ion setup (SBM 3), and all eight p-ions (SBM 8). Ee observed an increase in total identified PSMs by ∼7% (Fig. 4A). SBM 3 increased unique SUMOylated peptide IDs by 13% compared to the standard search and 20% compared to the VM search (Fig. 4B). Moreover, the data completeness increased from 44.1 to 45.3% when using SBM 3, indicative of more detection overall in 3/3 replicates. Intensity coverage (Fig. 4C), peak coverage (Fig. 4D), and the number of matched peaks, were all significantly increased with the addition of p-ions to the theoretical search space (Fig. 4E). When considering spectral annotation quality, we observed a notably increase for the SBM 3 search with 9% of the median Andromeda score (Fig. 4F). In general, increasing numbers of p-ions considered in the search resulted in larger numbers of matched peaks and correspondingly higher Andromeda scores of the SUMOylated peptides (Fig. S4). Importantly, scores for reversed database hits also increased with larger numbers of p-ions in the search space (Fig. S4D and Fig. S5D), stressing the need to carefully balance the number of included p-ions. Median PSM score remained identical for all non-SUMO target peptides (Fig. S5A). The median score decreased by 13.6 (20.1%) for the non-SUMO reverse hits (Fig. S5B). The median score increased by 6 (4.2%) for SUMO target hits (Fig. S5C) and by ∼1% for the SUMO reverse hits (Fig. S5D). Overall, the majority of all SUMOylated peptides were identified by all searches, with most unique identifications resulting from the SBM 3 and SBM 8 searches (Fig. 4G). The number of protein groups identified was 2884 for SBM 3, with 339 unique protein groups for this search, mainly identified from the nucleus (Fig. S6). The number of protein groups identified in SBM 8 was 2720, 2683 in Standard, and 2597 in the variable modification search (Fig. S6).

**Figure 3.**
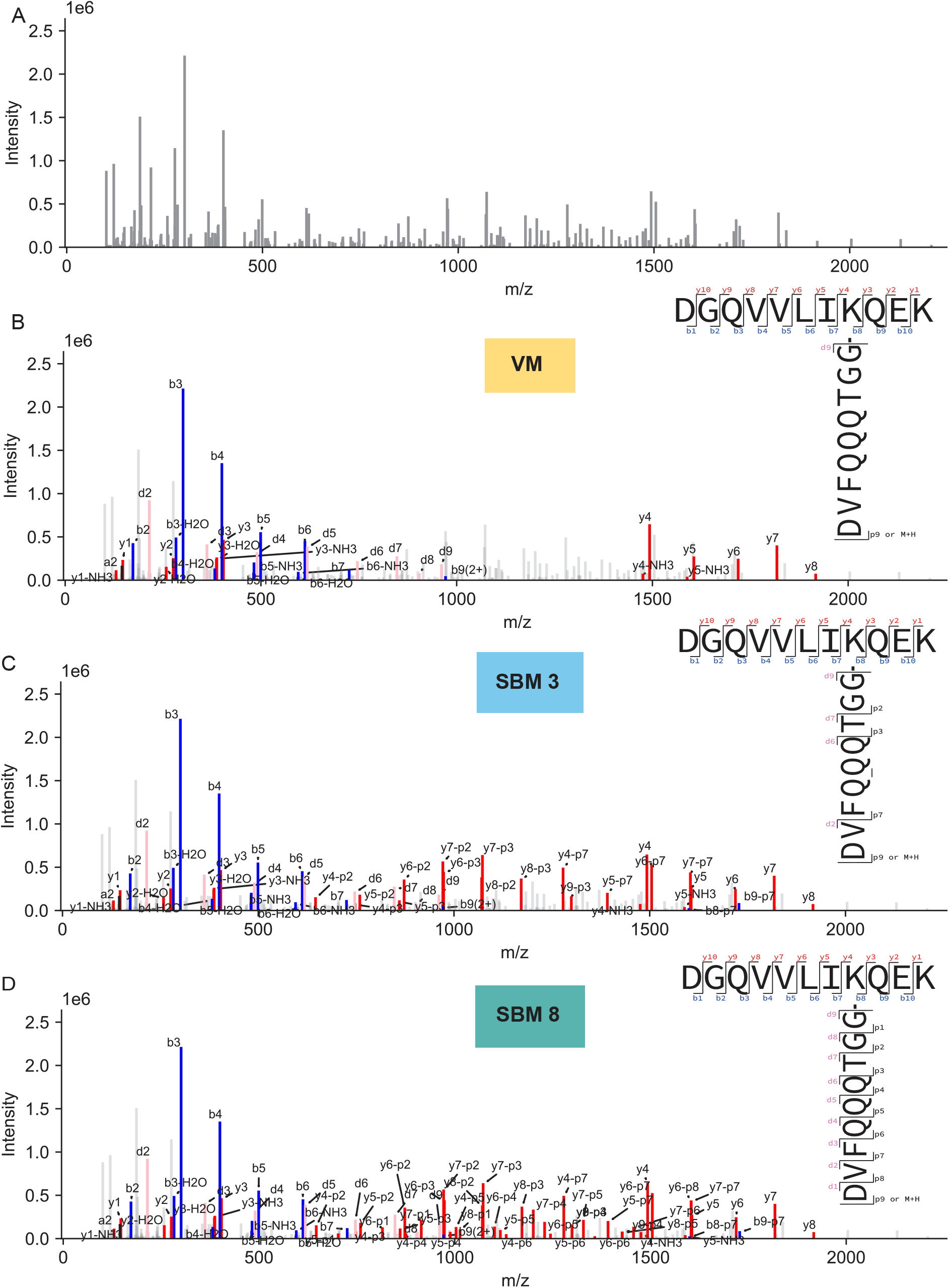
Spectrum of the identified SUMOylated target peptide DGQVVLIK(sumo)QEK, from the SBM 8 search (SUMO-HEK dataset). The top right SUMO peptide illustrations, coupled to each spectra, shows the theoretical fragmentations that were included in each search, generating the theoretical peaks for PSM generation. A. All mass deconvoluted m/z peaks of the selected spectrum, deisotoped and centroided. B. Highlighted annotations from the variable modification (VM) search space, where p-ions are not included, and SUMO is defined as a standard variable modification. C. Annotations from the SBM 3 search space, showing annotations for p2, p3, and p7 ions. D. Annotations from the SBM 8 search, displaying annotations for all included p-ions.

**Figure 4.**
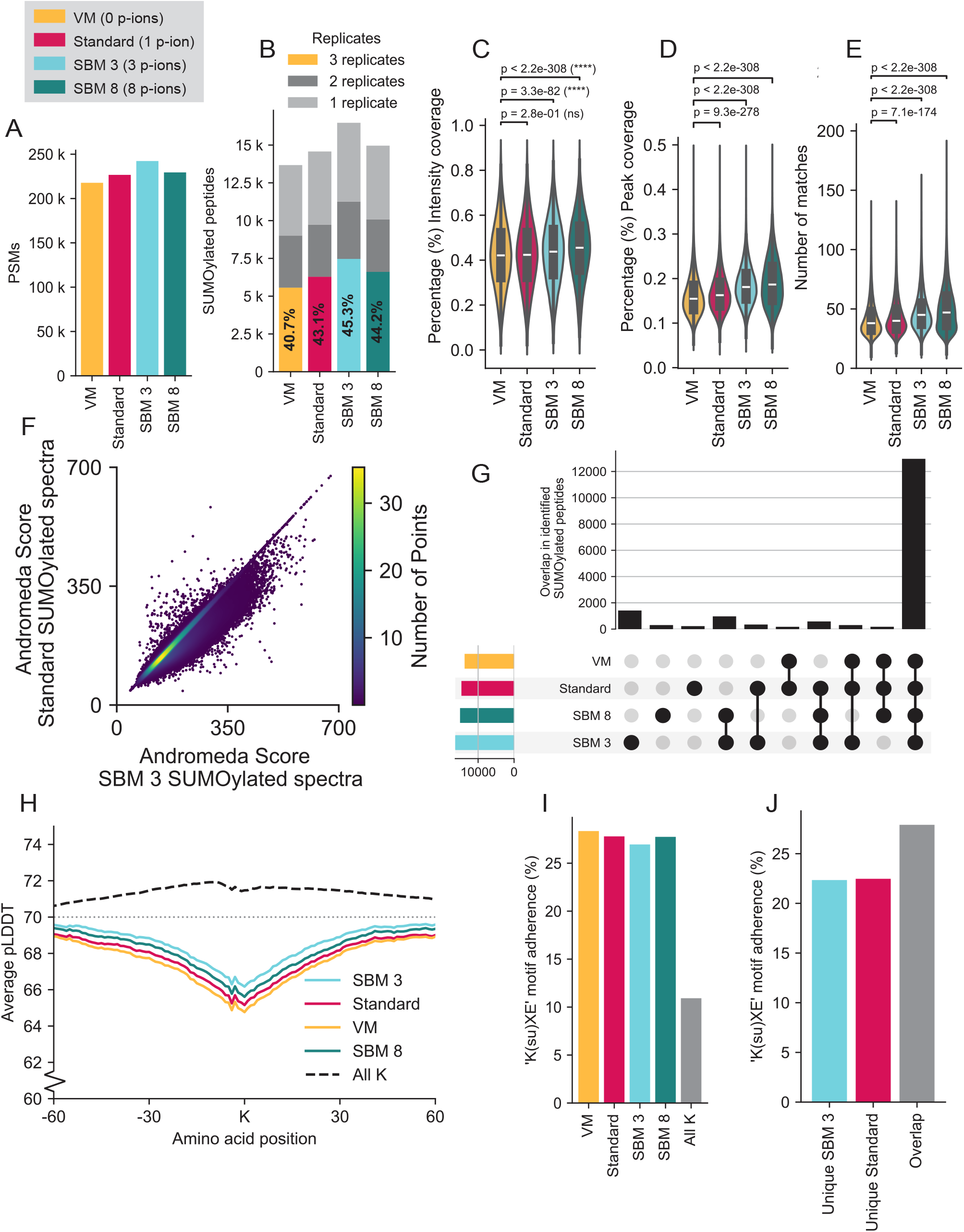
Comparison of the search result on analysing the SUMO-HEK dataset, with varying the number of p-ions in the searches: VM (0 p-ions), Standard (1 p-ion), SBM 3 (3 p-ions), or SBM 8 (8 p-ions). In all the figures, the reverse hits and contaminants have been removed. Violin plots illustrate the distribution of intensity coverage (%) across four datasets. Each violin includes an embedded boxplot showing the interquartile range, with the central line indicating the median. The number of data points (n) contributing to violin plots was 90,028 for VM, 97,699 for Standard, 112,409 for SBM 3, and 99,592 for SBM 8. A. The number of identified PSMs was 217745 for VM, 226671 for Standard, 242272 for SBM 3, and 229452 for SBM 8. B. Unique SUMOylated peptides identified across 1, 2, or all 3 replicates. C. Spectral intensity (abundance) coverage for the different searches. The median intensity coverage values for each dataset are as follows: VM (38%), Standard (40%), SBM 3 (45%), and SBM 8 (47%). D. Spectral peak coverage for the different searches. The median peak coverage values for each dataset are as follows: VM (15%), Standard (16%), SBM 3 (18%), and SBM 8 (19%). E. Average number of matched peaks for the different searches. The median number of matched peaks for each dataset is as follows: VM (38), Standard (40), SBM 3 (45), and SBM 8 (47). F. Scatter plot comparison of the Andromeda score between the SBM 3 search and the standard search. The average Andromeda score for SBM 3 is 200.8, while for standard it’s 183.9. G. UpSet plot visualizing unique and overlapping sets of peptides across the different searches. H. The average predicted local distance difference test (pLDDT) scores from AlphaFold-predicted protein structures are shown for regions flanking SUMOylation sites. Lower pLDDT scores correspond to regions with lower structural confidence, typically indicating intrinsic disorder. For comparison, pLDDT scores for regions surrounding all unmodified lysine residues (All K) in SUMO target proteins are also shown. The enrichment of low pLDDT scores near SUMO-modified lysines adheres to the notion that SUMOylation preferentially occurs in disordered regions [6]. I. KxE motif adherence for SUMOylated lysine residues across the different searches. The KxE motif adherence for unmodified lysine residues is shown for reference. J. KxE motif adherence across the overlapping SUMOylated peptides, as well as for search-specific peptides, comparing SBM 3 and standard.

### MaxSBM-exclusive SUMO sites conform to known biological properties

To validate SUMO sites identified in the SUMO-HEK dataset exclusively through the SBM module, we investigated two known biological properties of SUMOylation sites: the tendency for SUMO to reside in disordered protein regions [6] and adherence to the canonical KxE modification motif [6], [63]. To investigate SUMO-proximal structural properties, we extracted information from the AlphaFold structure database [60] and found that the pLDDT is lower near SUMOylated lysine residues compared to unmodified lysine residues (Fig. 4H). As a low pLDDT correlates with a higher degree of disorder, this validates the tendency for SUMO to occur in disordered regions. There were no notable differences in SUMO-proximal structural context between the different searches, suggesting that the SBM searches identify SUMO sites in similarly disordered regions. Adherence to the KxE consensus motif was 26.9% for SBM 3, 27.8% for standard, and 28.5% for the variable modification (VM) search (Fig. 4I). These numbers were considerably higher than the randomly expected occurrence of 11%, and overall agree with previous observations of KxE adherence for similar SUMOylome depth [6], [63]. For SUMO sites identified by both SBM 3 and standard search, KxE adherence was 27.9%, while peptides uniquely identified via SBM 3 or standard search were both ∼22.5%, indicating that neither search has a bias towards non-KxE motifs (Fig. 4J). Taken together, our observations support the notion that SUMO sites exclusively identified using MaxSBM adhere to known SUMOylation sequence preferences, and thus are not more likely to be false positives. Moreover, when considering SUMO target proteins, we found that these overall adhered to the subcellular localization expected for SUMO, e.g., predominantly nuclear and chromatin. SBM-exclusive SUMO target proteins showed a similar distribution and a preference to modify nuclear proteins, further substantiating the validity of the SBM search (Fig. S6).

### Comparison with existing tools for the identification of SBMs and labile PTMs

We established that MaxSBM improved performance for the detection of SUMO within the MaxQuant search engine. To benchmark the performance of MaxQuant against other software that can similarly handle SBMs, we utilized MSFragger-Labile [33] and compared performance on the SUMO-HEK dataset. Notably, MSFragger PTM output is post-processed using Percolator, and thus, we also adapted Percolator to post-process MaxQuant output generated using the SBM 3 search (Fig. 5). MSFragger-Labile with Percolator considerably outperformed MaxQuant in the absence of Percolator. Strikingly, when applying the Percolator pipeline to MaxQuant, the number of SUMO sites from the SBM 3 search was significantly increased by 70% (Fig. 5A), and slightly outperformed MSFragger with 13% more identifications. Notably, the majority of SUMO sites were found by all three approaches, followed by large clusters of sites identified by either or both of the Percolator-enhanced searches (Fig. 6B).

**Figure 5.**
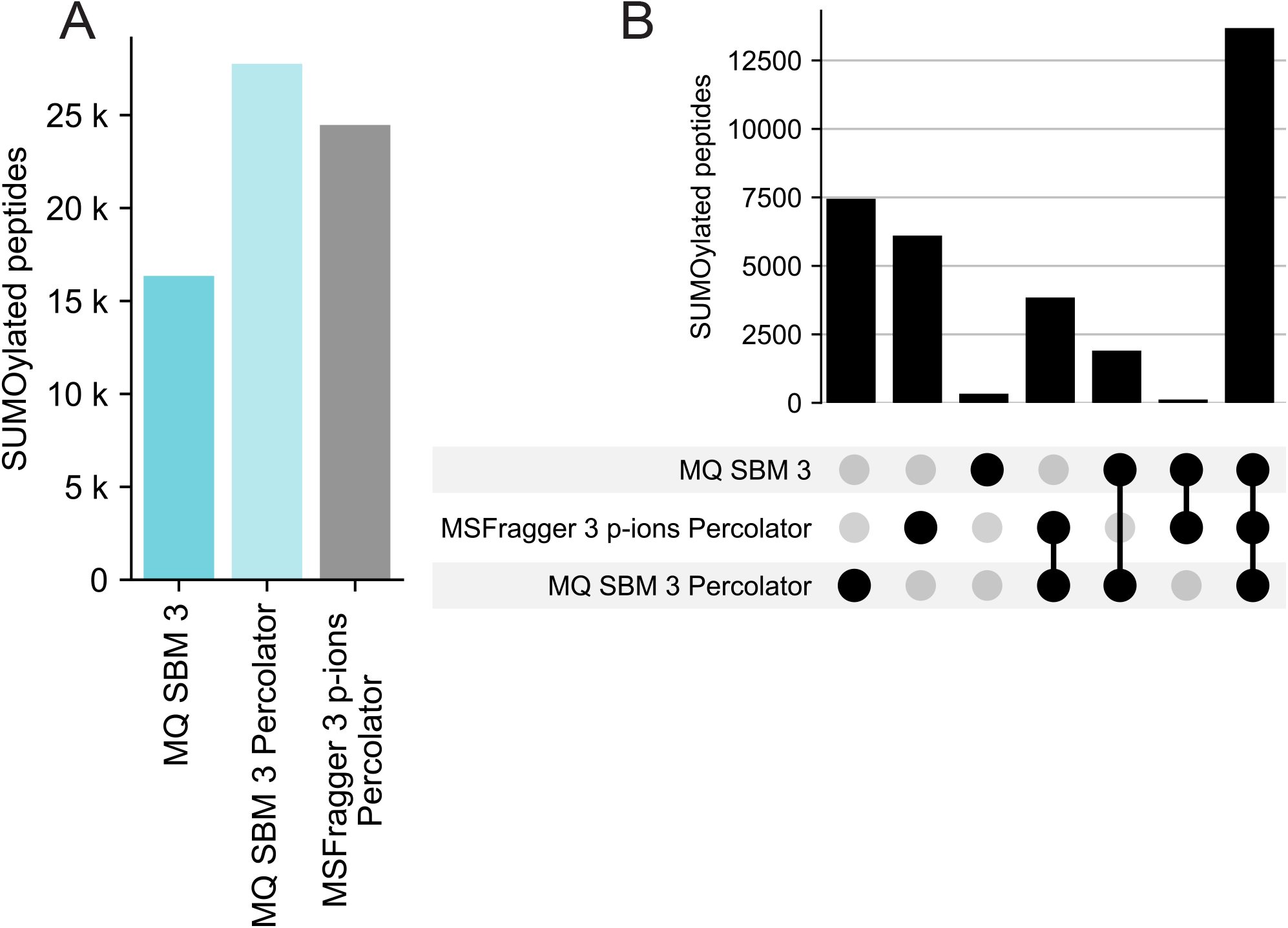
Comparison using existing tools on the SUMO-HEK dataset. A. The number of identified SUMOylated peptides with the SBM 3 search using MaxQuant FDR filtering was 16348, using Percolator for significance estimation gave 27762, and using MSFragger-Labile 3 p-ions gave 24469. All data was filtered to 1% FDR using the respective software options. B. UpSet plot visualization of unique and overlapping sets of SUMO sites, comparing MaxSBM 3, MaxSBM 3 with Percolator, and MSFragger-Labile with Percolator.

**Figure 6.**
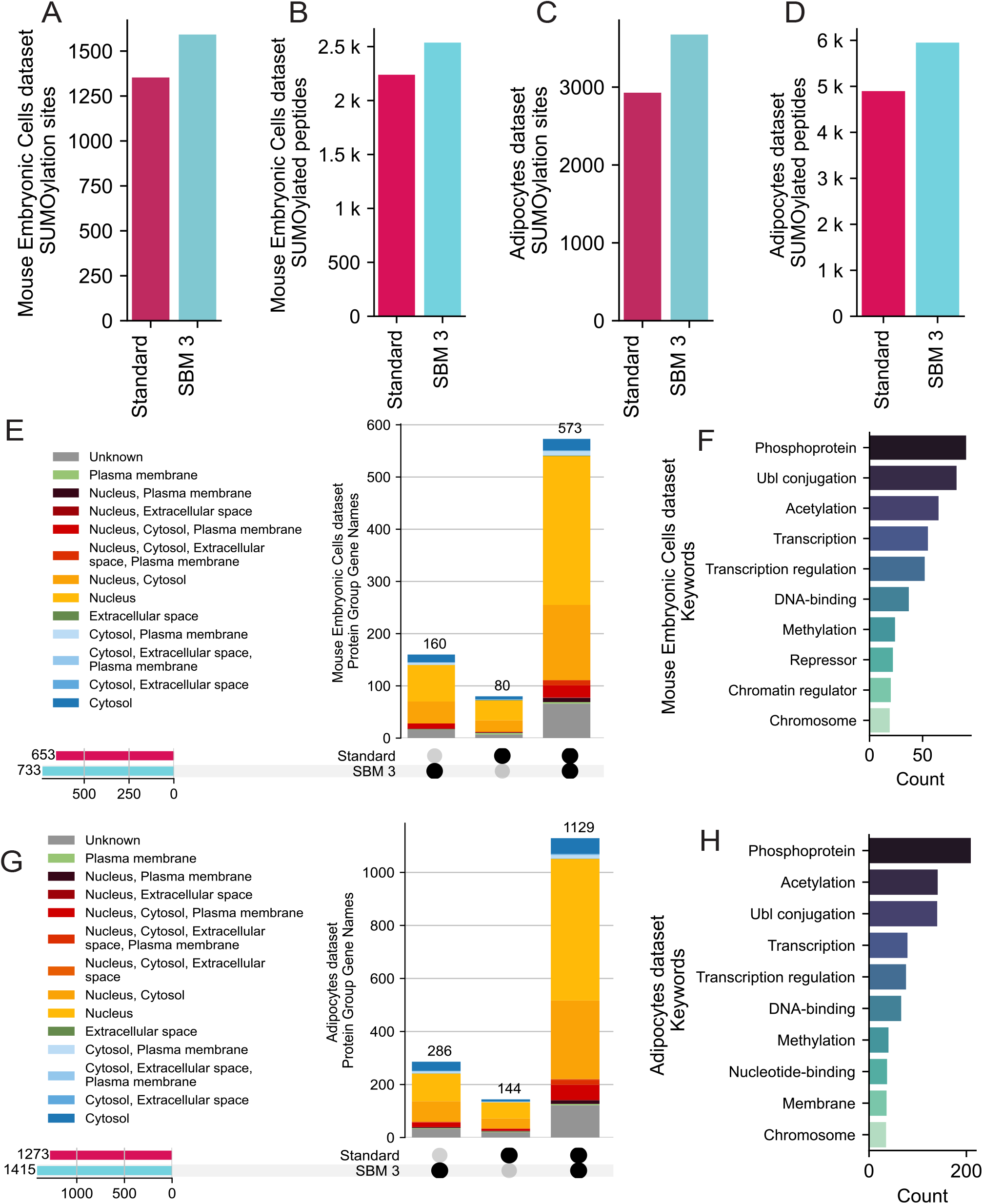
A. Number of identified SUMOylation sites in the SUMO-MEC dataset [19], derived from the ‘sites.txt’ MaxQuant output table. The optimized search SBM 3 is compared with a standard search. B. Number of identified SUMOylated peptides in the SUMO-MEC dataset, derived from the ‘msms.txt’ MaxQuant output table. C. Number of identified SUMOylation sites in the SUMO-Adip dataset [45]. D. Number of identified SUMOylated peptides in the SUMO-Adip dataset. E. The number of identified protein groups for the overlapping and unique sets in the SUMO-MEC dataset. The protein groups have been colored by their high-level cellular compartment [65]. F. Number of summed UniProt Keywords for the 160 unique proteins to SBM 3 in the SUMO-MEC dataset. G. The number of identified protein groups for the overlapping and unique sets in the SUMO-Adip dataset. H. Number of summed UniProt Keywords for the 160 unique proteins to SBM 3 in the SUMO-Adip dataset.

### Deriving Biological Insight

To further investigate the applicability of the MaxSBM module, we benchmarked the optimized search on two additional published endogenous SUMOylome datasets, with one based on mouse embryonic stem cells and fibroblasts (SUMO-MEC) [19], and the other based on differentiating mouse adipocytes (SUMO-HEK) [5], mouse embryonic cells (SUMO-MEC) [19], and mouse adipocytes (SUMO-Adip) [45]. In the embryonic stem cell dataset, we found an increase of 17.7% unique SUMOylation sites when using the module, from 1353 (Standard) to 1592 (SBM 3) (Fig. 6A). Importantly, the MaxSBM search profiled 160 SUMO target protein groups that were not found using a standard search, and reassuringly the SBM-exclusive proteins displayed a prominent nuclear localization (Fig. 6E). We investigated this subset of exclusive SUMO target proteins using the STRING database [64], and found them to be highly interconnected and displaying enriched biological terms that are expected from SUMO target proteins (Fig. S8), such as involvement in transcription and occurrence on phospoproteins (Fig. 6F). In the differentiating mouse adipocyte dataset, the module resulted in a striking 25.5% gain of SUMOylation sites (Fig. 6C), from 2928 (Standard) to 3675 (SBM 3). In addition, we found 286 SUMO target proteins not found using the standard search (Fig. 6G). Similarly, these exclusive proteins were primarily localized to the nucleus (Fig. 6G). Proteins specifically identified using the module were found to be highly interconnected using the STRING database (Fig. S9). Again, we found that these proteins were involved in DNA binding processes and transcriptional regulation (Fig. 6H). Altogether, by processing two physiological datasets using MaxSBM, we see substantial (18-26%) increases in the number of identified SUMO sites, along with overall higher spectral scores, and unveil hundreds of SUMO target proteins that align well with expected biological trends.

## Discussion and Conclusion

In this work, we demonstrate that inclusion of p-ions generated from SBMs (such as SUMO2/3) increases data completeness, peak coverage, and the intensity coverage, overall leading to a substantially better annotation and interpretation of existing data sets. To exemplify this, we used MaxSBM to re-process endogenous SUMO datasets, and can enhance peptide identification by 13% in a deep human cell line dataset [5], by expanding the theoretical peak list used for peptide score matching to include ions generated by fragmentation of the SUMO peptide remnant itself. We additionally show that we could increase the number of SUMOylation sites identified from mouse embryonic cells [19] by 17.7%, and from mouse adipocytes [45] by 25.5%. Importantly, the new sites derived via MaxSBM adhere to expected SUMO biological properties, including consensus motif adherence and structural preference. SUMO target proteins exclusively identified via MaxSBM reassuringly displayed predominant nuclear localization, as well as involvement in SUMO-related processes such as transcriptional regulation. Furthermore, the increased identification rate improves data completeness, i.e., detection of SUMOylation in more experimental replicates, which strengthens downstream quantitative proteomics analysis. Overall, MaxSBM enables an increased SUMO peptide and target protein coverage, and can be applied to previously recorded datasets, and may be beneficial for studying this pivotal post-translational modification.

Increasing the theoretical search space size is widely known in the field to decrease the identification rate [41], [42], [44]. While adding theoretical peaks will allow for more matches, this comes with a concomitant increase in the probability of a decoy match. Therefore, during a DDA database search, the search space should be kept as concise and specific as possible, while still encompassing all reasonably expected peptides [41]. Here, we find that the inclusion of the SBM p-ion series enhances both the sensitivity and specificity of SUMO peptide identification in database searches. However, the incorporation of the full set of eight p-ions resulted in fewer peptide identifications, likely due to highly increased odds of decoy matching. Our analysis indicated that the inclusion of three p-ions represented an optimal balance, which maximized identification performance without excessively inflating the search space. This is observed in both the highest identification rate in SBM 3 searches, as well as in the Andromeda score distributions. While the non-SUMO target hits remained the same, the SBM 3 module allowed for decreased scoring of non-SUMO reverse hits, enabling a better overall separation of the target-decoy distributions. In addition, the scoring increased for target SUMO hits in SBM 3, but interestingly not in SBM 8, showing that an additional search space increase was not beneficial. The observations are consistent with the well-established principle that expanding the number of theoretical fragment ions increases the probability of decoy matches, thereby reducing overall specificity. Consequently, while we here demonstrate the optimal p-ion assignment for the DVFQQQTGG peptide remnant, careful optimization would be essential for effective identification of other sequence-based modifications (SBMs) in data-dependent acquisition (DDA) workflows.

We logically also considered processing ubiquitin data sets following Lys-C digest, i.e., the UbiSite method [46]. However, after initial testing and benchmarking, we only saw very small increases (2%) in identifications using MaxSBM (Fig. S10). We reason that this could be due to the overall large size (∼1.4 kDa) of the ubiquitin Lys-C mass remnant, which contains multiple arginine residues, causing a very high (z=4-6) peptide charge, ultimately causing highly convoluted fragmentation spectra that remain a technical challenge to overcome.

Applying Percolator to estimate peptide significance yielded a marked increase in identification numbers for the endogenous HEK SUMO dataset. Specifically, the MaxSBM module alone resulted in a 13% increase in peptide identifications relative to the standard MaxQuant search, with improvements reaching up to 70% when Percolator was applied post-processing. This substantial gain was likely attributable to the inherent complexity of detecting SBM-modified peptides, resulting in limited score separation between target and decoy peptide-spectrum matches (PSMs). Percolator’s semi-supervised machine learning framework leverages additional features derived from PSMs, such as charge state, precursor mass error, and fragment ion coverage, to improve target-decoy discrimination. This capability could be particularly advantageous, where the difficulty in search space size limitations can cause scoring functions to underperform. However, despite the benefits, caution is warranted. While prior studies suggest that Percolator is relatively robust to overfitting due to its internal use of cross-validation and other regularization methods [44], [52], [66], the lack of an established ground truth in endogenous datasets poses a significant challenge for evaluating its performance objectively. However, other studies have presented examples where the FDR is not precisely controlled. Freestone et. al (2024, 2025) note that the validity of the cross-validation procedure might be compromised due to peptide species generating multiple spectra [67], [68], a problem that could be enhanced by the low signal in datasets during complex peptide identification. Therefore, the potential for Percolator to amplify the number of false positives cannot be entirely ruled out, particularly in cases where modifications introduce atypical and previously unseen fragmentation behaviors, which is particularly true for labile modifications and SBMs. In such contexts, thus, the impressive boost in identifications, while promising, underscores the need for rigorous validation. Future studies should further assess Percolator’s FDR estimates in the context of SBM detection. Finally, integrating emerging complementary tools, such as deep learning-based spectrum prediction, could further refine the discrimination between true and false identifications in modified peptide searches, although such approaches have yet to be implemented or validated in the context of sequence-based modifications.

In summary, the MaxSBM module notably enhances the identification of SUMOylated peptides, offering improved sensitivity and broader proteomic coverage in contemporary datasets, and could thus be expected to also enhance identification rates in future SBM datasets. By enabling more confident and comprehensive detection of modified peptides while maintaining robust statistics, MaxSBM facilitates downstream biological interpretation, including the mapping of modification-specific pathways, identification of modification hotspots, and characterization of dynamic regulatory networks. As such, this module represents a valuable tool for researchers seeking to elucidate the complex proteomic landscapes underpinning cellular signaling and post-translational regulation. Ultimately, advancements in this field can provide a foundation for incorporating spectral prediction and DIA into the analysis of SBMs.

## Abbreviations

SBM: (Sequence-Based Modifier)
SUMO: (Small Ubiquitin-like Modifier)
PTM: (Post-Translational Modification)
FDR: (False Discovery Rate)
PSM: (Peptide-Spectrum Match)

## Acknowledgments

We thank our colleagues at the Novo Nordisk Foundation Center for Protein Research and Max Planck Institute of Biochemistry for discussion and advice, in particular Nadezhda T. Doncheva and Jonas Damgaard Elsborg.

## Data availability

The module is integrated into MaxQuant search engine, which can be downloaded from www.maxquant.org [2025-02-07] [30], [40]. The software has been written in C# programming language using the.NET 8 framework. To run the application, Microsoft.NET 8 needs to be installed on the system. The software runs on Windows and Linux operating systems [69]. The Andromeda source code to calculate the theoretical SBM peaks can be found on GitHub: https://github.com/JurgenCox/mqtools7 [2025-06-18]. The scripts used for post-identification processing, diagnostic ion mining, protein inference, and msms.txt to Percolator input conversion can be found on GitHub: https://github.com/carolinelennartsson/SUMOylation [2025-06-18].

## Author Contributions

Conceptualization; CL, MLN and IAH. Data curation, investigation, and formal analysis; CL. Software development; CL and JC. Analysis validation; CL, PK, MLN, JVO, JC, and IAH. Visualization; CL and PK. Administration and supervision; MLN, JVO, JC, and IAH. First draft writing; CL. Final writing and editing; CL, PK, and IAH. All authors contributed to the article and approved the submitted version.

## Supplemental data

This article contains supplemental data, Supplemental Tables 1-2 and Figures 1-10.

