## Supplementary Information for "Improved peptide search for identification of SUMO and sequence-based modifications, in MaxSBM"

| **Sequence** | **d-ion** | **d-ion Composition** | **d-ion neutral Mass (Da)** | **Corresponding p-ion** |
| --- | --- | --- | --- | --- |
| E | d1 | H(7) C(5) N(1) O(3) | 129.04 | p12 |
| ES | d2 | H(12) C(8) N(2) O(5) | 216.07 | p11 |
| EST | d3 | H(19) C(12) N(3) O(7) | 318.12 | p10 |
| ESTL | d4 | H(30) C(18) N(4) O(8) | 431.21 | p9 |
| ESTLH | d5 | H(37) C(24) N(7) O(9) | 567.27 | p8 |
| ESTLHL | d6 | H(48) C(30) N(8) O(10) | 680.34 | p7 |
| ESTLHLV | d7 | H(57) C(35) N(9) O(11) | 779.41 | p6 |
| ESTLHLVL | d8 | H(68) C(41) N(10) O(12) | 892.50 | p5 |
| ESTLHLVLR | d9 | H(80) C(47) N(14) O(13) | 1048.61 | p4 |
| ESTLHLVLRL | d10 | H(91) C(53) N(15) O(14) | 1161.68 | p3 |
| ESTLHLVLRLR | d11 | H(103) C(59) N(19) O(15) | 1318.79 | p2 |
| ESTLHLVLRLRG | d12 | H(106) C(61) N(20) O(16) | 1375.81 | p1 |
| ESTLHLVLRLRGG | d13 | H(109) C(63) N(21) O(17) | 1431.83 | - |

**Supplemental Table 1.** Chemical compositions and monoisotopic neutral masses of Ubiquitin d-ions and corresponding p-ions.

| **Input Feature** | **Type** | **Description** | **MaxQuant Column(-s)** |
| --- | --- | --- | --- |
| SpecId | String | Composite spectrum identifier with decoy/target tag | Reverse, Scan number, Raw file, Charge |
| Label | Integer | Label used for Percolator target/decoy classification | Reverse |
| ScanNr | Integer | Scan number from the original spectrum | Scan number |
| ExpMass | Float | Experimental precursor m/z value | m/z |
| CalcMass | Float | Calculated monoisotopic mass [M+H]+ | Mass |
| lnrSp | Float | Natural log of the ranking based on the Score | Logarithm of ranked index based on Score |
| deltLCn | Float | The difference between this PSM's Score and the Score of the last-ranked PSM for this spectrum, divided by this PSM's Score or 1, whichever is larger. | Score - Last score / (Score or 1, if Score < 1) |
| deltCn | Float | The difference between this PSM's Score and the Score of the next-ranked PSM for this spectrum, divided by this PSM's Score or 1, whichever is larger. | Delta score / (Score or 1, if Score < 1) |
| Sp | Float | Score | Score |
| IonFrac | Float | Fraction of matched fragment ions | Peak coverage |
| Mass | Float | Calculated peptide mass | Mass |
| PepLen | Integer | Length of peptide in amino acids | Length |
| ChargeN | Integer | Precursor charge state | Charge |
| enzInt | Integer | Number of missed cleavage sites | Missed cleavages |
| enzN | Boolean | Whether peptide has enzymatic N-terminus (based on cleavage rules) | Calculated based on enzymatic clevage pattern |
| enzC | Boolean | Whether peptide has enzymatic C-terminus (based on cleavage rules) | Calculated based on enzymatic clevage pattern |
| Peptide | String | Modified peptide sequence | Modified sequence |
| Proteins | String | Protein accession(s) matched to the peptide | Proteins |

**Supplemental Table 2.** The Percolator input features from, summarized from the CRUX documentation page: <https://crux.ms/file-formats/features.html> [2024-08-15]. The table contains the input features with their description and the corresponding MaxQuant column. A python script for the full calculations and conversation from MaxQuant msms.txt is available in the GitHub repository: <https://github.com/carolinelennartsson/SUMOylation> [2024-08-15].

A.

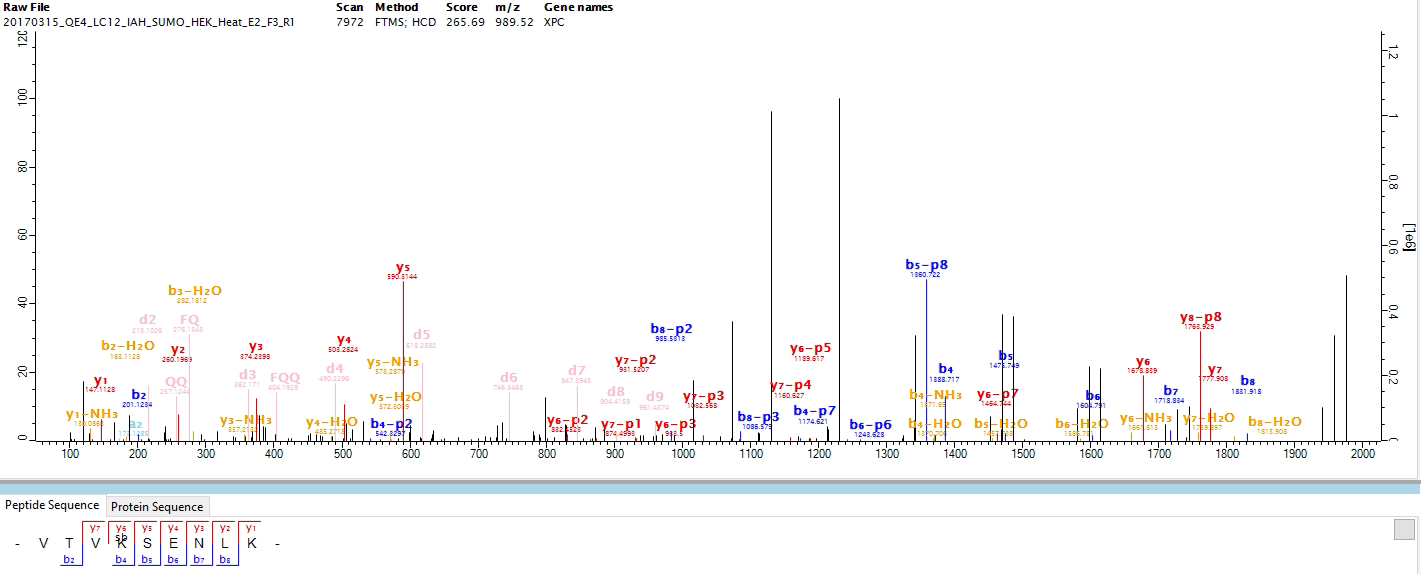

B.

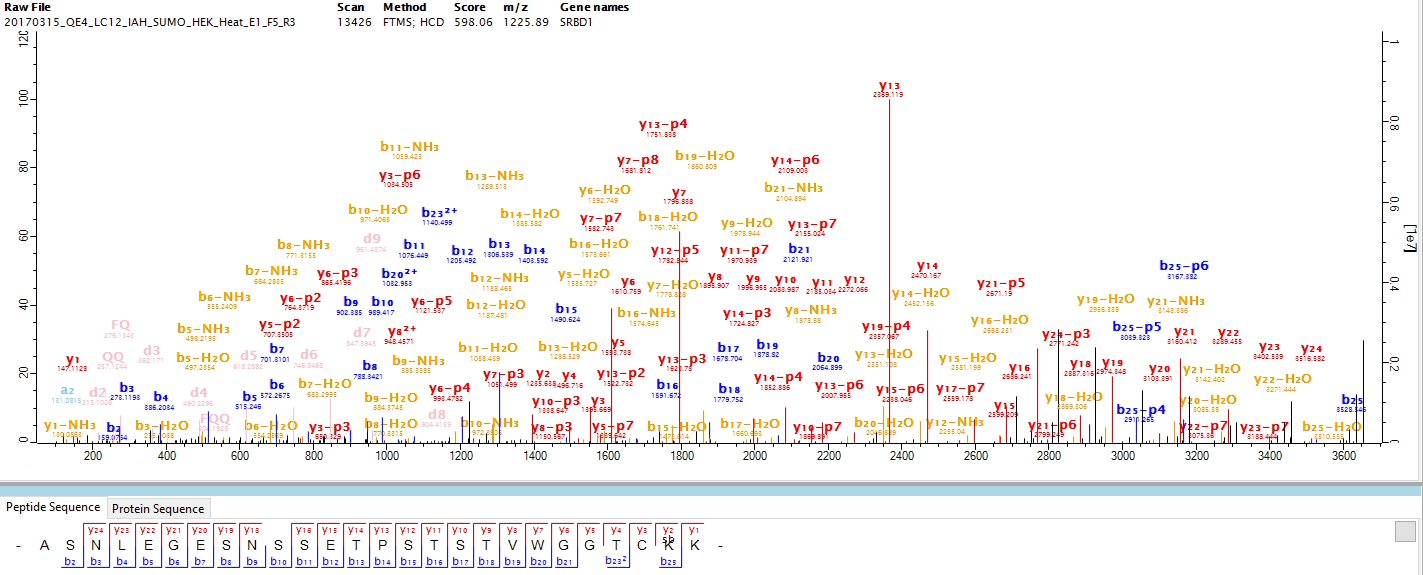

**Supplemental Figure 1.** **A and B.** Spectral view of identified SUMOylated peptides inside the MaxQuant Viewer [1]. The y-ion are annotated in red, the b-ions in blue, the d-ions in pink and molecule loss ions in yellow. The p-ions that are introduced in this module are shown with their respective b and y annotation, in their respective colour.

A.

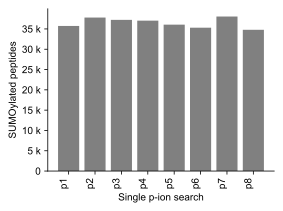

B.

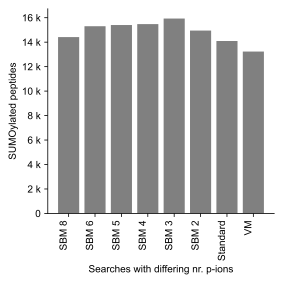

**Supplemental Figure 2.  A**. Search using a single p-ion to detect SUMOylated peptides in the endogenous HEK dataset [2]. This approach tests the sensitivity of minimal ion evidence by restricting the search to only one fragment ion characteristic of SUMOylation, serving as a baseline for comparison with more complex ion combinations in B. **B.** Optimization of the MaxSBM search on a subset of the endogenous dataset containing SUMOylated peptides. The optimization tested combinations of 0 (VM), 1 (Standard), 2, 3, 4, 5, 6, and 8 p-ions. Results were compared to a standard search using SUMO as a variable modification (VM).

A.	B.

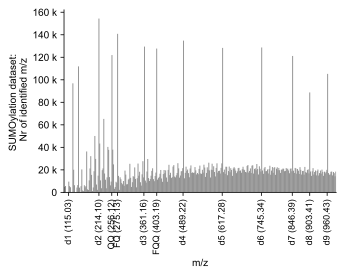

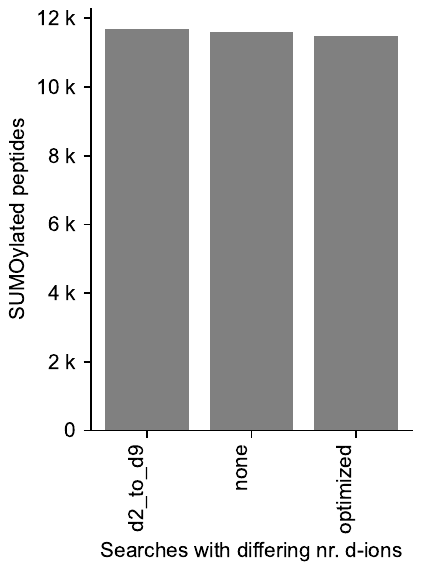

C.	D.	E.

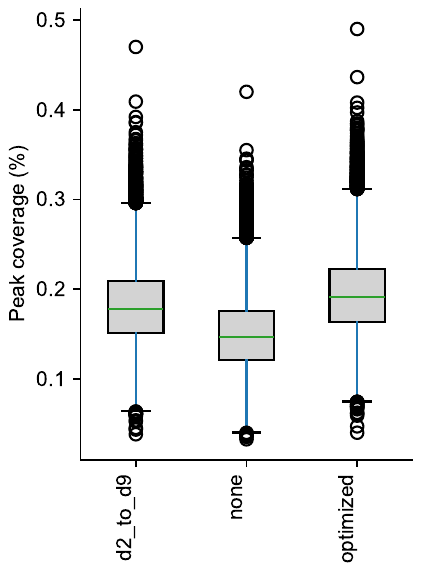

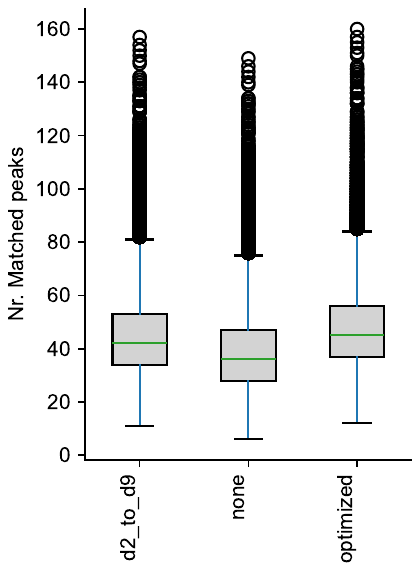

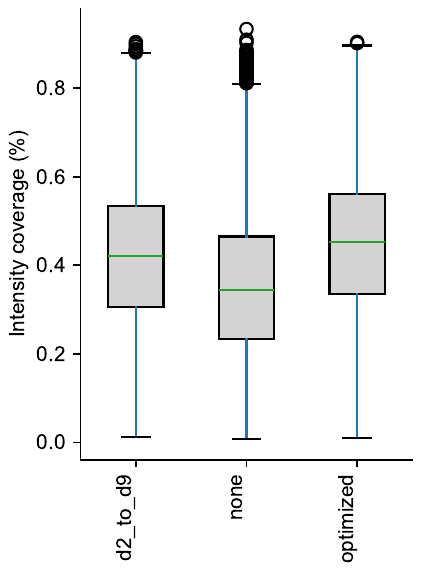

**Supplemental Figure 3. A.** Diagnostic ion mining of SUMOylation d-ion peaks. All masses across the endogenous HEK dataset [2] were summed to observe the commonly measured peaks, revealing the d-ion series, including the masses corresponding to the sequences ‘FQQ’, ‘QQ’, and ‘FQ’. The figure was based on the data from ‘msms.txt’ from the SBM 3 search. **B**. Identified a unique number of SUMOylated peptides. Result from d-ion optimization, from a subset of the HEK endogenous dataset [2]. In optimization search “d2_to_d9”, d-ions d2 to d9 were included. In search ‘none’, none of the d-ion were included, and in “optimized” search, d2 to d9 were included, with the addition of the masses corresponding to the double cleaved SUMO remnant corresponding to the sequences: ‘FQ’, ‘QQ’, and ‘QQQ’. **C.** Peak coverage of the d-ion optimization searches. **D.** Number of matches peaks of the d-ion optimization searches. **E.** Intensity coverage of the d-ion optimization searches.

A. B.

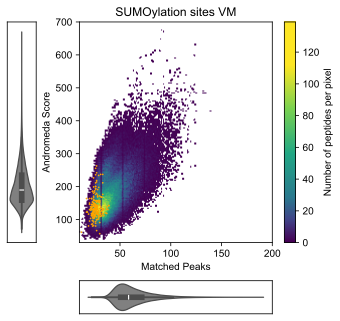

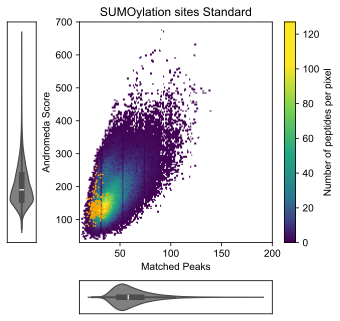

C. D.

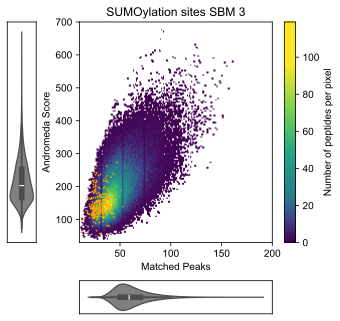

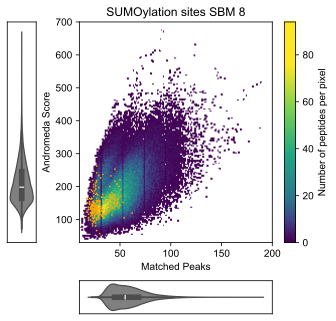

**Supplemental Figure 4.** Searches comparing the andromeda score and number of matched peaks for different number of p-ions in the endogenous HEK dataset [2]. The reverse hits are shown in orange. **A.** Search with SUMOylation as a variable modification (VM). Median score: 168.4; median matched peaks: 38. **B.** Standard search with one p-ion included. Median score: 169.3; median matched peaks: 40. **C.** Search with 3 p-ions included (SBM 3). Median score: 182.3; median matched peaks: 45. **D.** Search with 8 p-ions included (SBM 8). Median score: 177.1; median matched peaks: 47.

A.

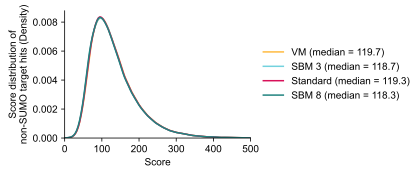

B.

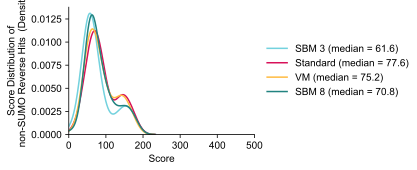

C.

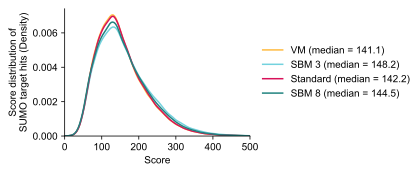

D.

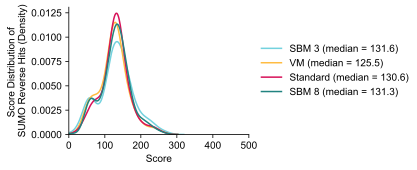

**Supplemental Figure 5.** Andromeda score distributions of the target and decoy hits, from the endogenous HEK dataset [2]. **A.** Score distribution of all non-SUMO target PSMs, yielding a 0.6 difference (<<1%) in median score, comparing SBM 3 to Standard. **B.** Score distribution of all non-SUMO reverse hits, yielding a 13.6 decrease in median score (~20.1%), comparing SBM 3 to Standard. **C.** Score distribution of SUMOylated target hits, yielding a 6 increase in median score (~4.2%), comparing SBM 3 to Standard. **D.** Score distribution of all SUMOylated reverse hits, yielding 1 median score increase (~1%), comparing SBM 3 to Standard.

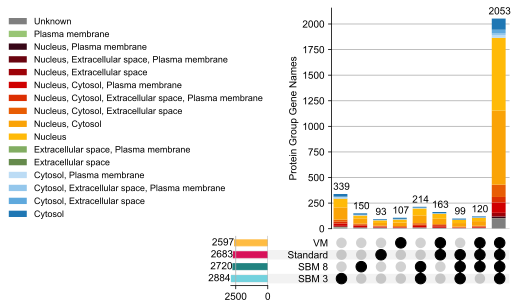

**Supplemental Figure 6.** Identified SUMOylated proteins in the endogenous HEK dataset [2], comparing the different searches SBM 3, 8, VM and Standard. The protein groups have been coloured by their high-level cellular compartment, derived from the COMPARTMENTS database [3].

A. B.

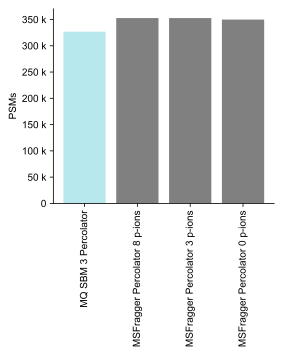

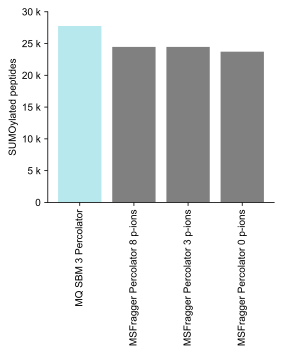

**Supplemental Figure 7.** MSFragger-Labile [4] searches comparing MaxQuant SBM 3 search (with Percolator used to estimate significance) with MSFragger-Labile hybrid mode, from the endogenous HEK dataset [2]. MSFragger was ran with 0, 3 and 8 p-ions as with Percolator as postprocessor. **A.** Number of PSMs identified with different number of remainder ions included in MSFragger-Labile. **B.**  Number of identified SUMOylated peptides identified with different number of remainder ions included in MSFragger-Labile.

A.

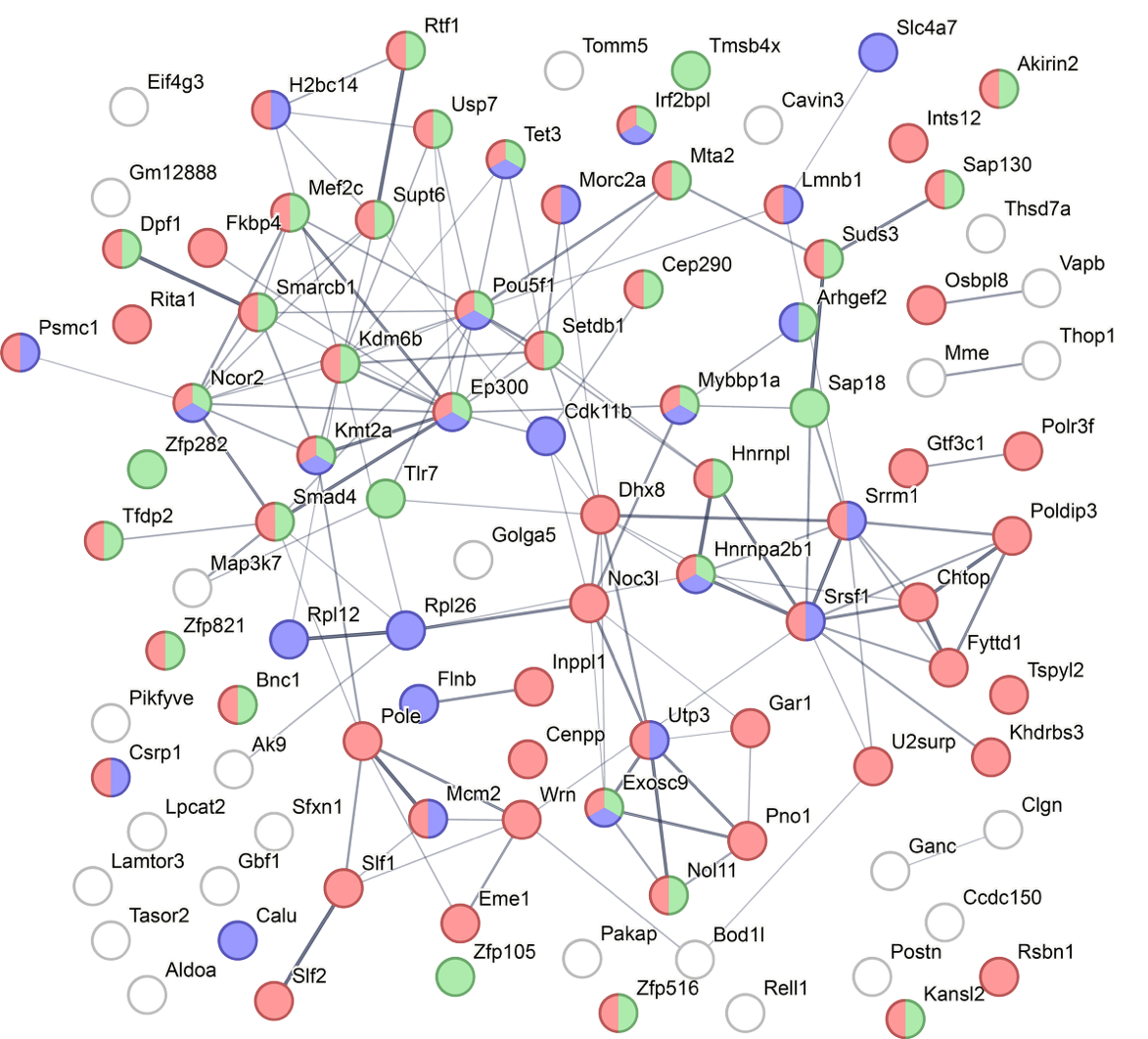

B.

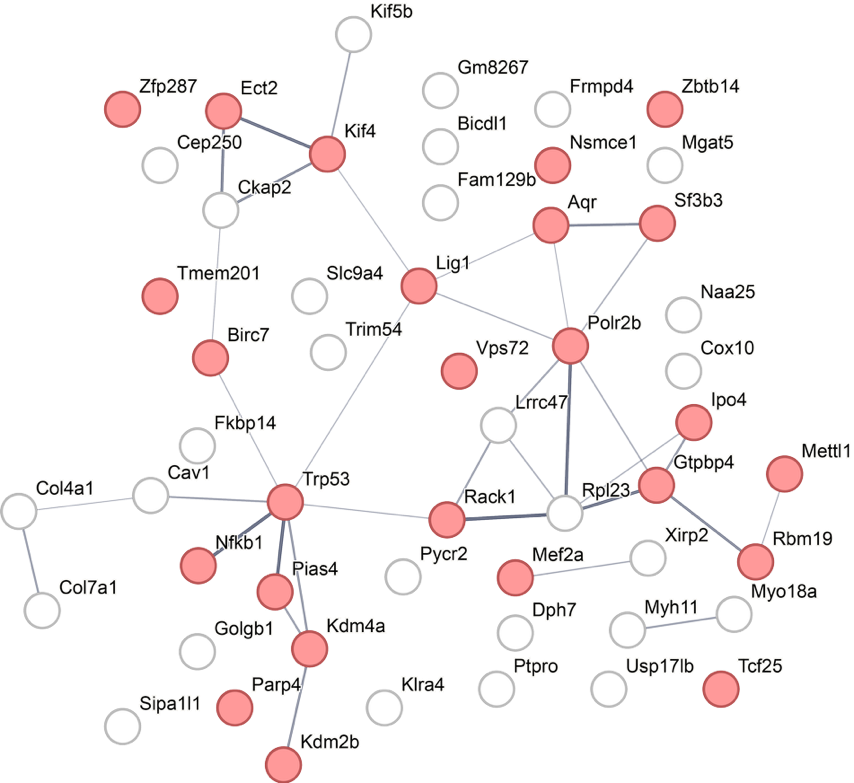

**Supplemental Figure 8.**  STRING [5] protein interaction network with proteins identified with the module, in the mouse embryonic cells dataset. Coloured by enriched terms; red = nucleus, blue = embryo tissue expression, and green = regulation of transcription. **A.** The proteins unique to the SBM3 search: <https://version-12-0.string-db.org/cgi/network?networkId=bXy42HHpDwUK> **B.** Proteins uniquely identified to standard: <https://version-12-0.string-db.org/cgi/network?networkId=bmEvqomUYpcN>

A.

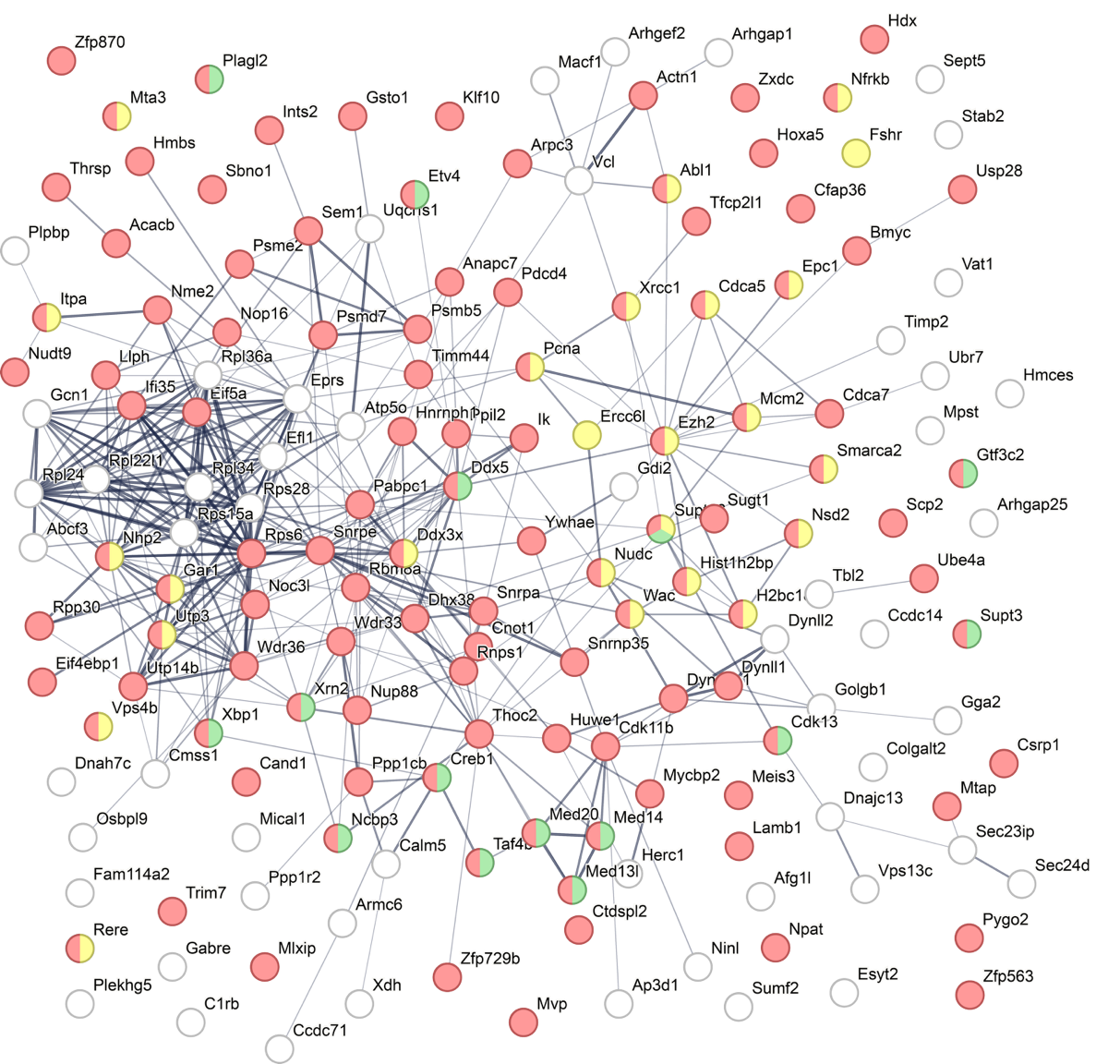

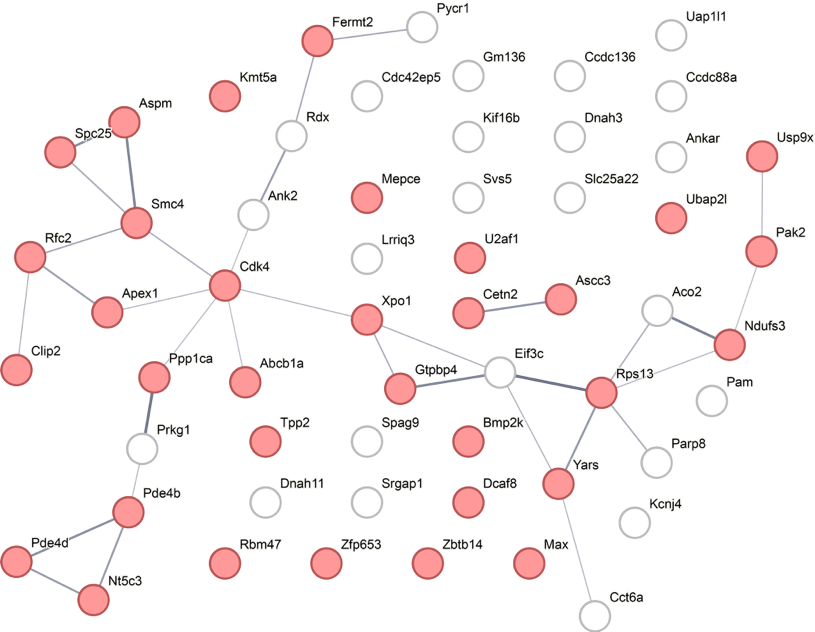

**Supplemental Figure 9.** STRING [5] protein interaction network with proteins identified with the module, in the mouse adipocytes dataset. Coloured by enriched terms; red = nucleus, yellow = chromosome organisation, green = regulation of transcription **A.** The proteins unique to the SBM3 search: <https://version-12-0.string-db.org/cgi/network?networkId=bF9mFzB1SnNl> **B.** Proteins uniquely identified to standard: <https://version-12-0.string-db.org/cgi/network?networkId=bEskDKsOingt>

A. B.

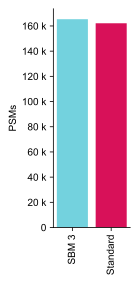

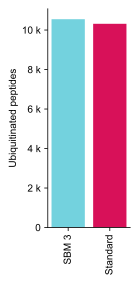

**Supplemental Figure 10.** Primary analysis runs on the ubiquitin dataset, generated with the UbiSite method [6] (PXD006201). The samples were digested and run with Lys-C enzyme, different from the standard protocol that uses trypsin. **A.** Number of total identified PSMs, comparing the SBM 3 search with a standard search. **B** Ubiquitin identified ubiquitinated peptides, 10548 for SBM 3 and 10316 for Standard search. Yielding a 2% increase in unique ubiquitinated peptides.
